# Spatial chemistry of citrus reveals molecules bactericidal to *Candidatus* Liberibacter asiaticus

**DOI:** 10.1101/2024.04.12.589303

**Authors:** Alexander A. Aksenov, Alex Blacutt, Nichole Ginnan, Philippe E. Rolshausen, Alexey V. Melnik, Ali Lotfi, Emily C. Gentry, Manikandan Ramasamy, Cristal Zuniga, Karsten Zengler, Kranthi Mandadi, Pieter C. Dorrestein, Caroline Roper

## Abstract

Huanglongbing (HLB), associated with the psyllid-vectored phloem-limited bacterium, *Candidatus* Liberibacter asiaticus *(C*Las), is a disease threat to all citrus production worldwide. Currently, there are no sustainable curative or prophylactic treatments available. In this study, we utilized mass spectrometry (MS)-based metabolomics in combination with 3D molecular mapping to visualize complex chemistries within plant tissues to explore how these chemistries change *in vivo* in HLB-impacted trees. We demonstrate how spatial information from molecular maps of branches and single leaves yields insight into the biology not accessible otherwise. In particular, we found evidence that flavonoid biosynthesis is disrupted in HLB-impacted trees, and an increase in the polyamine, feruloylputrescine, is highly correlated with an increase in disease severity. Based on mechanistic details revealed by these molecular maps, followed by metabolic modeling, we formulated and tested the hypothesis that *C*Las infection either directly or indirectly converts the precursor compound, ferulic acid, to feruloylputrescine to suppress the antimicrobial effects of ferulic acid and biosynthetically downstream flavonoids. Using *in vitro* bioassays, we demonstrated that ferulic acid and bioflavonoids are indeed highly bactericidal to *C*Las, with the activity on par with a reference antibiotic, oxytetracycline, recently approved for HLB management. We propose these compounds should be evaluated as therapeutics alternatives to the antibiotics for HLB treatment. Overall, the utilized 3D metabolic mapping approach provides a promising methodological framework to identify pathogen-specific inhibitory compounds *in planta* for potential prophylactic or therapeutic applications.

## Introduction

Huanglongbing (HLB) is a highly destructive citrus disease exhibiting complex symptomatology. All commercial varieties are HLB-susceptible to various degrees, posing a global threat to production^1^. HLB is caused by the bacterium *Candidatus* Liberibacter asiaticus (*C*Las) and is spread by the insect vector *Diaphorina citri,* commonly known as the Asian citrus psyllid (ACP). Early HLB symptoms include leaf mottling, yellow shoots and rapid tree death^2^. Moreover, the effect on fruit development has major downstream consequences on fresh fruit and juice quality. Fruit from HLB-impacted trees are small, misshapen, with irregular maturation patterns that yield unpalatable flavor profiles.

The ACP, first reported in Florida in 1998, catalyzed rapid regional HLB outbreaks by 2005, resulting in endemic establishment and billions in industry damages^31^. ACP has since spread east to west across the US citrus belt, and HLB is now detected in thousands of Californian urban citrus plants, threatening commercial groves^4^. Disease management has mainly focused on insect vector control with repeated sprays of synthetic insecticides^5^, and no sustainable bacterium-targeting HLB interventions yet exist.

Currently, oxytetracycline antibiotic trunk injection is the primary approach against *C*Las^6,7,8^. However, antibiotic deployment is an unsustainable strategy that engenders both human health and environmental risks, given the propensity for bacterial resistance. A “precautionary tale” - widespread antibiotic administration has led to the rapid emergence of resistant *Candidatus* Liberibacter asiaticus strains in China^9^. Thus, alternative bactericidal agents need to be explored for viable and sustainable approaches to *C*Las treatment.

Metabolomics, both nuclear magnetic resonance (NMR) and mass spectrometry (MS)-based, gives valuable insights into molecular distributions, including those underlying pathogenesis, and thus may provide leads on potential therapeutic molecules. Past investigations employing NMR and MS established associations between *C*Las infection, such as terpenoid/sugar disruption underlying symptomatic flavor decline^10^. However, uneven HLB distribution hinders generalizing systemic changes^11,12^. As symptoms manifest locally, pinpointing locally perturbed metabolites is essential to understand the disease and propose targeted therapies.

Recent advances in metabolomics enable exploring spatial chemical distributions^13–16^. We hypothesized that detailed infection mapping would expedite discovery of compounds active against *C*Las *in planta*, to inform novel HLB therapeutics. In this work, we visualized the HLB effect on the metabolome of citrus plants using 2D and 3D molecular maps to gain pathological insights, illuminating possible bacterium-mediated host metabolite exploitation. While focused on citrus, this approach, in principle, should be generally applicable to a wide range of plant-pathogen systems.

## RESULTS

### Metabolome shifts in field and greenhouse trees with HLB

We carried out an untargeted metabolomics analysis of citrus tree tissues across a spectrum of disease severity collected from seven HLB-infected Florida orchards, where the disease is endemic. Tissues throughout the citrus tree: stems, roots, and leaves were collected and analyzed. Because of the complex metabolome of citrus trees across different tissues, we used molecular networking^17,18^ to map and explore the detected chemistries. The resulting network grouped molecules that fragmented in a similar fashion, thus capturing their structural similarity and aiding in understanding the chemical distributions^18,19^. The network of the detected metabolome across grove trees is shown in Figure 1 a,b. The metabolome varied as HLB progressed (Figure 1c), with molecules associated with disease severity forming distinct clusters in the network. This suggests that metabolic disruption in HLB progresses not through individual molecular changes, but shifts across entire biochemical families. Clusters of compounds - dilinolenins, fatty amides like octadecenamide, glycerophosphocholines, terpenoids, and, prominently, flavonoids - showed changes linked to symptom severity across tissues. To control for confounding factors such as length of infection, local conditions etc., samples were also collected from a controlled greenhouse environment, including infected and uninfected healthy trees (confirmed using qPCR). Metabolome differences related to HLB symptomatology were much more pronounced in greenhouse samples as compared to field samples. The most discerning diagnostic markers, identified via partial least squares discriminant analysis (PLSDA) and Random Forest, were found to be various flavonoids. This implicates the flavonoid pathway as the main metabolomic “barometer” of infection^11,20–22,23,24–26^.

**Figure 1.**
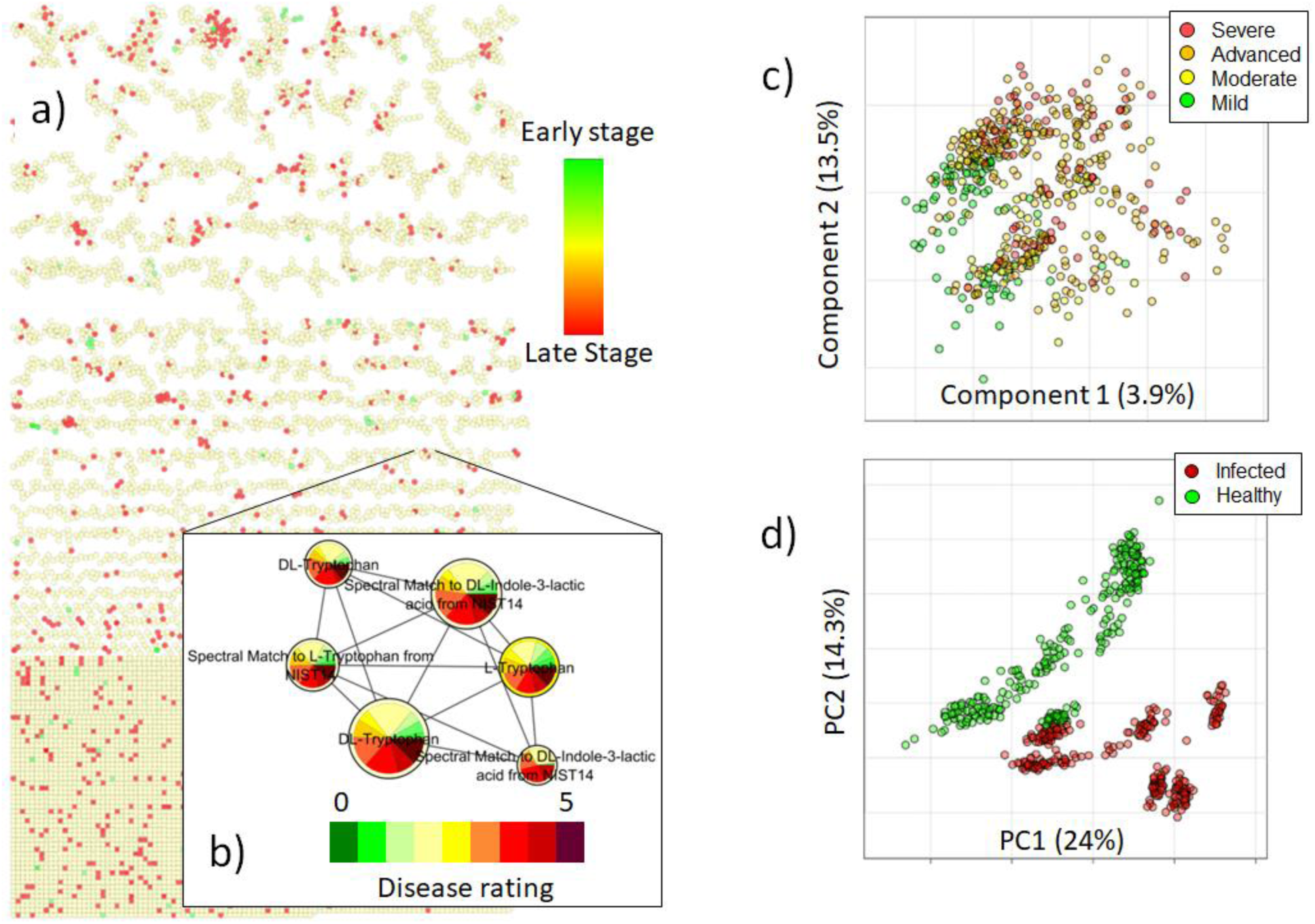
Global network analysis of metabolomic shifts in HLB-impacted trees. Citrus trees (leaves, stems and roots) were sampled across multiple groves in Florida. a) Global network of the metabolome detected in samples collected from grove trees in Florida, which included trees exhibiting symptoms across a range of disease severity (1 -appears healthy to 5 - severe HLB symptoms). b) A close-up of a network cluster showing distributions of molecular abundances in trees with different disease ratings (higher values correspond to higher symptom severity); node size is related to total compound abundance. The compounds in the cluster were present in higher amounts when the disease was more severe. The selected example shows perturbation in amounts of tryptophan, indicative of altered metabolism, and a metabolite known to be of microbial origin, indole-3-lactic acid (ILA), a tryptophan metabolite that is known to play a role in microbe-host interactions; ILA is associated with increased disease severity. c) A supervised analysis (partial least squares discriminant analysis, PLSDA) of tissues from trees in orchards across Florida showed metabolome stratification according to disease severity. The analysis indicates differences in metabolomes associated with disease severity (Q2 0.1775). d) Unsupervised analysis (principal component analysis, PCA) of tissues from greenhouse-reared trees indicated drastic differences in the metabolome of healthy and infected symptomatic tissues.

### 3D Metabolome Mapping

To explore the spatial patterns in distribution of metabolites, molecular maps were generated using the ‘ili tool ^13^. We further elucidated these patterns for HLB-discriminating molecules. This mapping was conducted on tissues from greenhouse-reared trees of the same age and length of infection. 3D metabolome maps were rendered by collecting individual leaves along an infected branch and mapping specific metabolite concentrations, such as the flavonoid scutellarein tetramethyl ether, on the to branch images (Fig. S1). This allowed for visualization of metabolite concentration along the same branch where apical leaves exhibited HLB symptoms and basal leaves are asymptomatic (Fig. S1a). Notably, the distribution of the flavonoid scutellarein tetramethyl ether may be related to leaf age rather than symptomology. Therefore, the symptom-related patterns were investigated at single-leaf resolution, where leaves are age-matched. At this scale, symptom-related patterns were prominent, with various flavonoids sharing similar spatial propagation pattern; depleted in visibly chlorotic and mottle signatures of infection (Fig S2). At the same time, oxidized flavonoids tended to exhibit opposite spatial patterns, suggestive of mutual interconversion (Figure 2, S3).

**Figure 2.**
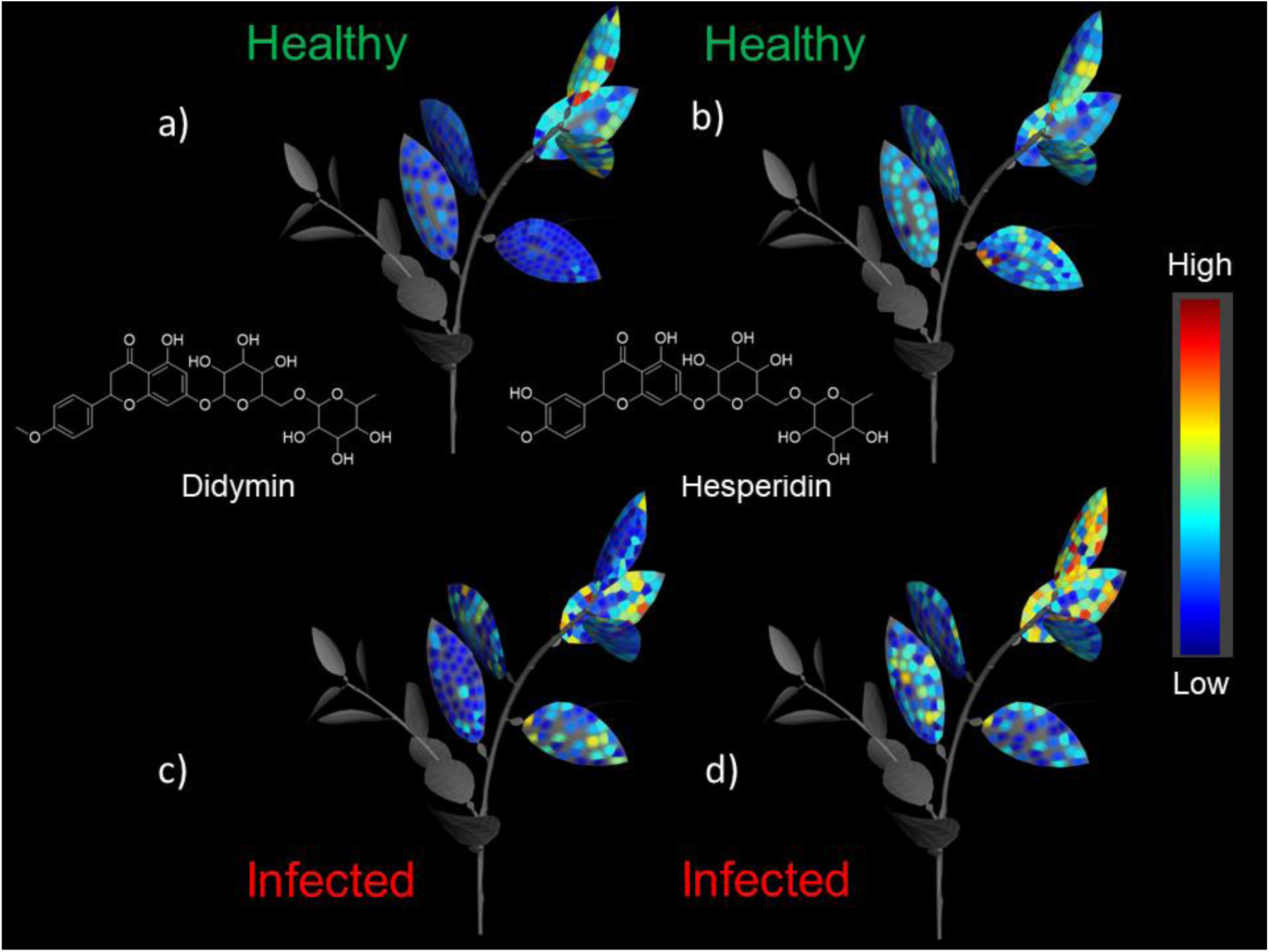
3D mapping of the distribution of the flavonoids, didymin and hesperidin in infected and healthy citrus branches: a) Didymin (m/z 595.2026 (rt 208.16)), b) Hesperidin m/z 611.1973 (rt 131.25)) and infected plant: c) Didymin and d) Hesperidin. Didymin was present at higher abundance in younger leaves of healthy plants and was depleted in the infected plant, while hesperidin increased in abundance in the infected plant and didymin was depleted, indicating possible formation of hesperidin due to oxidation of didymin.

Other than flavonoids, we noted feruloylputrescine, a conjugate of ferulic acid (part of the flavonoid biosynthesis pathway) and putrescine as molecules discriminating of HLB symptoms. Feruloylputrescine is preferentially accumulated in apical symptomatic leaves, exactly opposite the non-oxidized flavonoid depletion patterns (Fig S4a). The ferulic acid abundance decreased concomitantly with putrescine conjugation, indicative of the possible conversion into feruloylputrescine (Fig S4b). The same trends were evident on the single-leaf scale, with a clear correspondence of spatial distributions to the chlorotic spots phenotype (Figure 3). Another related phenolic acid, hydroxycinnamic acid (*p*-coumaric acid), showed a similar distribution, enriched in symptomatic areas (Fig S5, 3c).

**Figure 3.**
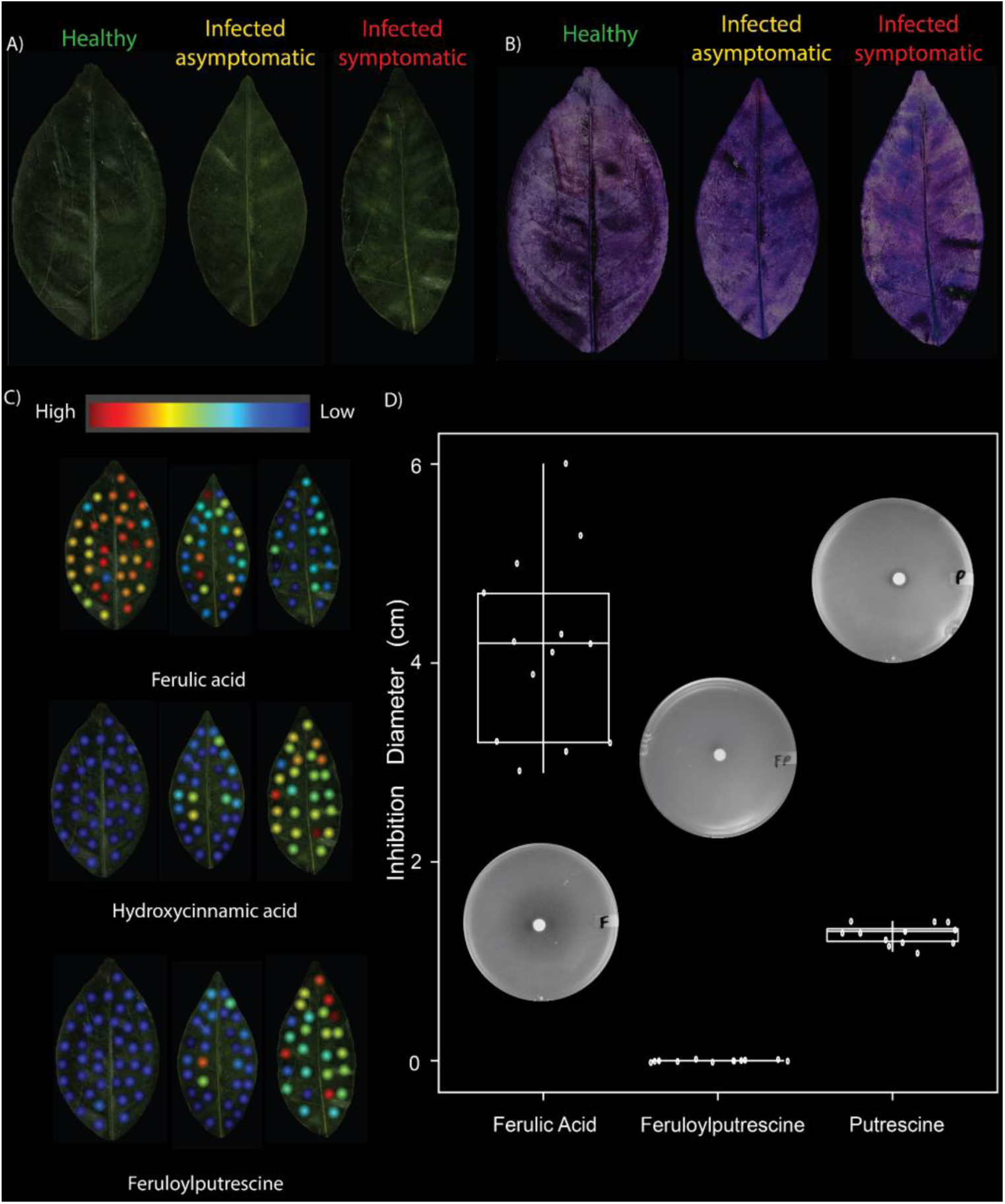
2D mapping of polyamine derivatives within single leaves. a) Images of leaves with different disease severity; b) The leaves shown in (a) in a color scheme that clearly shows chlorotic spots (deep blue shade areas) c) Mapped distributions of (top to bottom): ferulic acid - H_2_O (only water loss ion was observed), hydroxycinnamic acid and feruloylputrescine. 2D maps: healthy, infected asymptomatic, infected symptomatic (from left to right). The mapped symptomatic and asymptomatic leaves were sampled from two branches of a tree that was confirmed by qPCR to be infected with *C*Las (C_t_ = 19.6). Increases in hydroxycinnamic acid and feruloylputrescine were concomitant with increased disease severity, while ferulic acid exhibited the opposite trend. d) Plot showing *Liberibacter crescens* (a culturable surrogate for *C*Las) growth inhibition assay, when challenged with 35 mg/mL solution by ferulic acid, feruloylputrescine or putrescine as quantified by the diameter of the zone of inhibition. Ferulic acid had a pronounced inhibition effect.

### Metabolic modeling of *C*Las uptake rates of plant metabolites

A possible rationale for the observed spatial patterns in the metabolome (in particular, the low abundance of ferulic acid in symptomatic tissue) may be the metabolism of ferulic acid by the bacterium that would protect itself from antimicrobial activity. A supporting argument for this would be the observation that phenolic acids, like ferulic acid, reduce the growth rate of *C*Las, while in conjugation with putrescine, this effect is reduced due to the synthesis of feruloylputrescine. Therefore, we tested this hypothesis by conducting model simulations using the previously established genome-scale metabolic network of *C*Las^9^. We simulated uptake rates of plant metabolites by *C*Las for compounds that are part of arginine and proline metabolism, as well as those present in the biosynthesis of phenylpropanoids. The model predicted that uptake rates of ferulic acid between 1×10^-10^ and 1×10^-8^ mmol/gDW/h would be sufficient to reduce the growth rate of *C*Las up to 90%. These predictions were experimentally confirmed and putrescine uptake rates in this range did not predict a reduction of *C*Las growth. Interestingly, when these ferulic acid uptake rates (1×10^-10^ and 1×10^-8^ mmol/gDW/h) were combined with putrescine uptake rates between 1×10^-2^ and 10×10^-1^ mmol/gDW/h, the model predicted the highest *C*Las proliferation (Figure 4).

**Figure 4.**
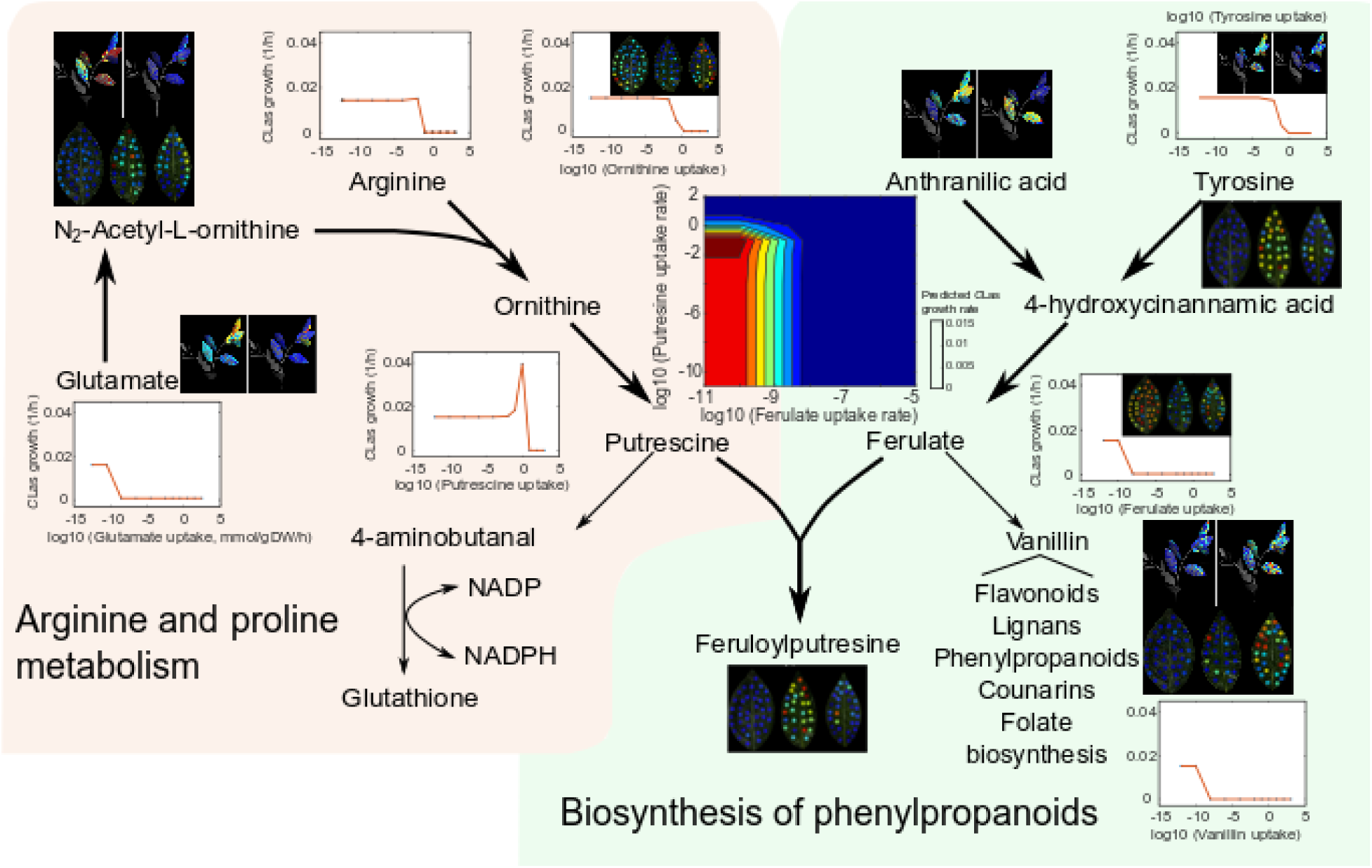
Genome-scale metabolic network simulations about the effect of feruloylputrescine biosynthesis on the growth of *C*Las. Contour plot shows the predicted growth rates of *C*Las, while varying putrescine and ferulate uptake rates. Individual scatter plots show the response of *C*Las growth to changes in the individual uptake rates of metabolites of interest (mmol/gDW/h). 2D maps of leaves show from left to right: healthy, infected symptomatic, infected asymptomatic samples of citrus plants. 2D maps of plants show from left to right: healthy and infected citrus plants.

### *In vitro* verification of model predictions with disc assay

In order to further confirm *in silico* predictions, we assayed putrescine, ferulic acid, and feruloylputrescine for potential antibacterial activity using a previously developed disc diffusion assay based on *Liberibacter crescens*, a culturable surrogate of the unculturable *C*Las^27^. Metabolic modeling of the pathogen described in the previous section predicted putrescine and feruloylputrescine to be mildly beneficial and ferulic acid to be inhibitory to *C*Las growth. The model predictions were consistent with the metabolites’ spatial distributions in greenhouse citrus tree leaves. Disc assay results showed that putrescine and ferulic acid were indeed moderately and highly inhibitory to *L. crescens,* respectively. The conjugated metabolite, feruloylputrescine, was not inhibitory to *L. crescens* growth as predicted by our model (Figure 3d). These *in vitro* results, in conjunction with the observed interconversion between ferulic acid, putrescine, and feruloylputrescine in HLB symptomatic trees, indicate that *C*Las likely directly, or through manipulation of host metabolic pathways, conjugate ferulic acid to putrescine producing the non-toxic feruloylputrescine, which increasing its survivability in the host.

### *Ex vivo* verification of model predictions with *C*Las-citrus hairy root assay

To further validate the toxicity of ferulic acid and flavonoids, whose biosynthesis is disrupted due to conjugation, against the target pathogen *C*Las, we conducted a *C*Las-citrus hairy root cultures assay^28^. As a possible low-cost, accessible therapeutic intervention, we tested over-the-counter citrus peel extracts as a widely available source of native citrus bioflavonoids. *C*Las-containing hairy roots were treated with bioflavonoids and ferulic acid for 72 h (Figure 5a). Untreated and ethanol (0.2% v/v) samples were used as negative controls. A reference antibiotic, oxytetracycline (OXY), reported as an inhibitor of *C*Las^298,30^, was used as a positive control. After treatment, all tissue samples were exposed to PMAxx dye (propidium monoazide, Biotium, Fremont, CA) that allows measurement of only live *C*Las bacterial DNA, following DNA extraction and molecular diagnostics. The relative titers of *C*Las were estimated using qPCR. We found that bioflavonoids significantly inhibited *C*Las (p ≤ 0.05) in a dose-dependent manner (125, 250, and 500 ppm), whereas ferulic acid showed inhibition at 125 and 250 ppm (p ≤ 0.05), on par with oxytetracycline when compared to untreated controls (Figure 5b).

**Figure 5.**
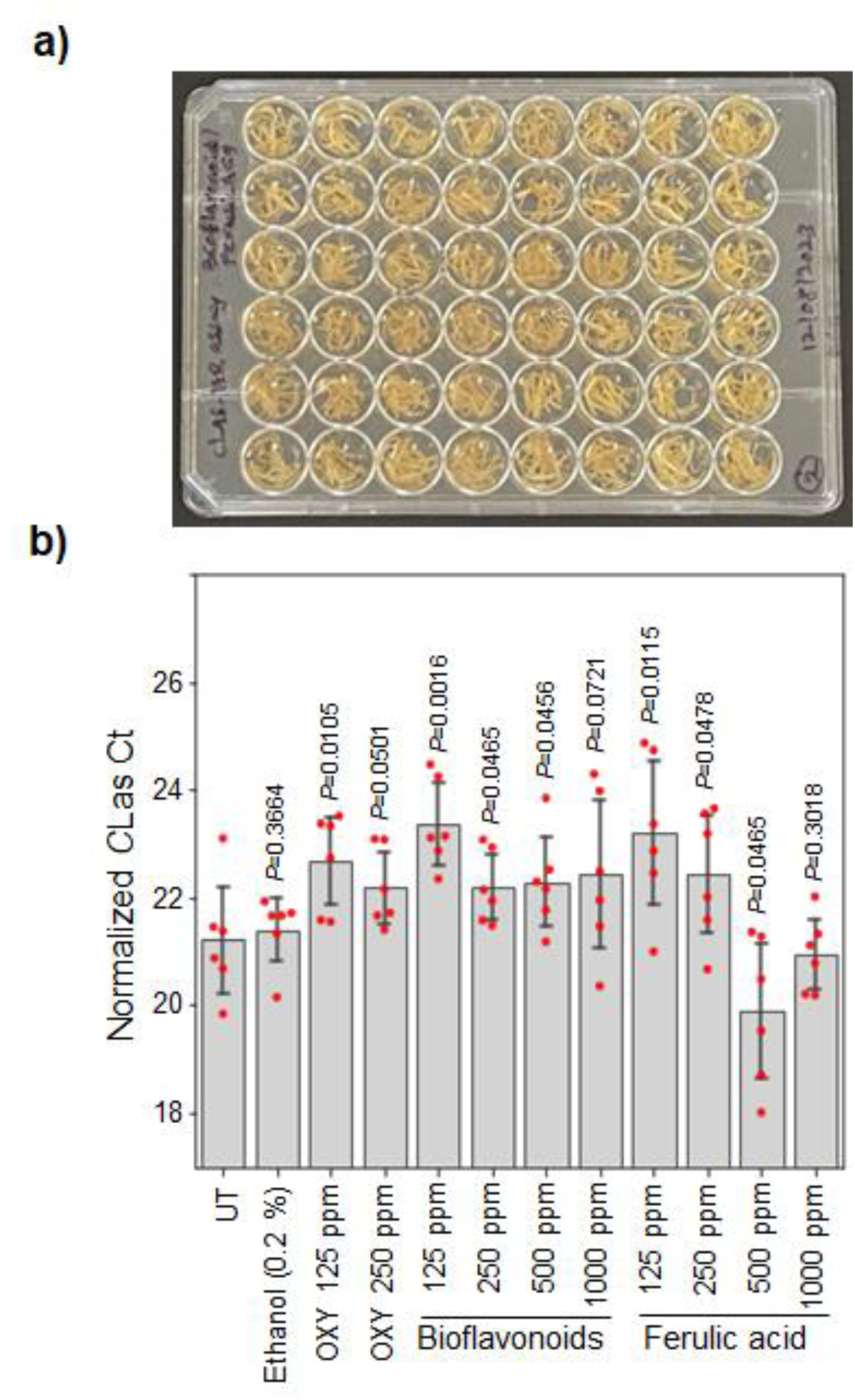
Bioflavonoids and ferulic acid inhibit the growth of *Candidatus* Liberibacter asiaticus (*C*Las) in *C*Las-citrus hairy roots. (a) The *C*Las-citrus hairy roots were treated for 72 h with 125, 250, 500, and 1000 ppm of bioflavonoids and ferulic acid. Oxytetracycline hydrochloride (OXY)-treated hairy roots (250 and 500 ppm) were used as a positive control, and untreated (UT) ethanol (0.2%) used to dissolve the compounds was used as a negative control. Relative titers of *C*Las (b) were estimated after 72 h of treatment, followed by qPCR analysis. Error bars represent the standard error of the mean (n=6), and p values were calculated by Student t-test relative to untreated samples.

## DISCUSSION

We explored HLB disease metabolome dynamics using spatial mapping, predictive modeling, *in vitro,* and *ex vivo* assays to identify endogenous citrus compounds with plant therapeutic potential. Metabolites altered by HLB symptoms were consistent among field and greenhouse trees, with flavonoids comprising the most discriminating “biomarkers”, supporting previous observation^11,31,32,30^. The discriminant flavonoids structures did not comport to a notable trend, and molecules differing by various backbone substitutions, e.g. addition of a glycan (which increases the compound’s solubility), were found to be predominantly declining in infected symptomatic tissues. This implies upstream flavonoid pathway suppression that potentially aids pathogen establishment, fitting the known antimicrobial and defense roles of these compounds. However, this information alone is insufficient to understand disease etiology^31,33,34,30^. For example, certain flavonoids were instead found to increase with HLB symptom development.

Overlaying 3D distribution onto molecular networking provided further insights into functional interconnections in molecular distributions, not accessible otherwise. For example, the association with HLB symptoms is notably different for oxidized versions of some flavonoids. As shown in Figure 2, hesperidin accumulation mirrored declines in the flavonoid didymin, possibly reflecting oxidative conversion of the latter into the former during infection. Such localized oxidative shifts were further evident across flavonoid families (Figure S3), aligning with oxidative stress reports previously described^35,36^. In another example, 3D maps revealed that leaf age was the predominant driver of flavonoid abundance, rather than disease severity. Younger leaves possess elevated flavonoids, necessitating age-matching to correctly determine infection biomarkers.

Exploring the HLB-discriminating ability and corresponding 3D maps for various metabolites allowed us to identify other molecules of interest in addition to flavonoids. In particular, feruloylputrescine was noted to be highly associated with symptom severity. Distribution patterns implicated the conversion of ferulic acid into feruloylputrescine, mirrored by ferulate disappearance and putrescine buildup (Figure 3). This shift was further echoed by hydroxycinnamate rises, likely reflecting the oxidation of cinnamic acid precursors. Taken together, spatial patterns indicate metabolic channeling away from phenylpropanoid biosynthesis toward polyamine conjugation due to infection.

Exploring upstream pathways points to a branch point: while general phenylalanine supply appeared unaffected, p-coumaroyl-CoA intermediates depleting from the flavonoid pathway were instead visible as hydroxycinnamate en route to feruloylputrescine. This suggests that *C*Las has evolved to counteract both ferulic acid and flavonoids formation, ostensibly as a self-defense against their toxicity.

To confirm that these molecules are indeed toxic to *C*Las, we conducted the disc and hairy roots assays that confirmed their bactericidal effects toward *C*Las. Although the disc assay exploited a surrogate organism, *Liberibacter crescens,* the hairy roots assay is a way to directly assess the toxicity, both genetic- and chemical-based to *C*Las itself ^28,37,38^. In this approach, *C*Las is maintained in citrus hairy root tissues, and the antimicrobials are directly infiltrated into the roots by vacuum, thus overcoming the delivery bottleneck that exists with whole tree assays^7,29,39^ and circumventing concomitant irreproducibility issues. Additionally, because citrus hairy roots have intact vasculature, they are much closer to the *in planta* citrus environment where *C*Las resides, thus providing reliability over testing against surrogate bacteria or heterologous systems^40,41,42,43^. The assay has demonstrated a clear bactericidal effect of the flavonoids on par with the anti-*C*Las activity of oxytetracycline. In the case of Ferulic acid, the lower dosages (125 to 250 ppm) are highly effective, while increasing the dosage to 500 or 1000 ppm did not seem to be effective, possibly due to the onset of toxicity of the compounds to the hairy roots themselves^44^. Both flavonoids and Ferulic acid bacterial inhibitory abilities have been well characterized in other systems, but the mode of action is not yet established^45,46^.

These phytochemicals may disrupt bacterial cell-to-cell communication^47^, modify biofilm production and inhibit swimming motility^48^, impact bacterial membranes^49^, or even create pores or rupture cell members^50^. Nevertheless, these experiments confirmed the robust anti-*C*Las activity of the ferulate and flavonoids^50,49^. It is not known how these compounds may impact other members of the citrus microbiome. Indeed, both bacterial and fungi communities drastically shift as HLB symptoms advance^34^, and citrus-associated fungi can produce *Liberibacter* spp. inhibitor compounds^27^. Future studies should consider the interactions of these compounds with the microbiome, and vice versa, to increase efficacy of disease management applications.

With the efficacy against *C*Las confirmed, the ferulic acid and bioflavonoids present an opportunity for field application as novel Huanglongbing therapies. These compounds are plant-derived, non-toxic, biodegradable, inexpensive to manufacture at scale, and thus would be a very appealing class of therapeutics. They may help to bypass negative environmental and economic externalities compared to current methods, such as antibiotic use and synthetic chemical pesticide applications for ACP control^6–9^. While optimal therapy delivery systems for citrus plants remain an active topic of research and development, multiple practical options such as trunk injection already exist^51–53,34,7,8,30,54^.

The impact of HLB has been devastating. Deployed alongside psyllid population suppression, the proposed “botanical” solutions harnessing natural plant defenses, when deployed at scale, may offer a viable path towards stable HLB mitigation and control. Finally, although we demonstrate the described approach for the citrus and *C*Las, the same approach of mapping the metabolic distributions coupled with toxicity assays may present opportunities to generalize the development of a therapy agent discovery pipeline that could be used for a wide range of pathosystems.

## Materials and Methods

### Tissue collection and processing

#### Branch and single leaf assay

For the branch and single leaf assays, Valencia sweet orange (*Citrus sinensis*) grafted onto Swingle rootstock was propagated in a greenhouse and inoculated with *C*Las using infectious ACP. Five ACP adults enclosed in a nylon mesh drawstring bag were applied and confined to newly emerged leaves for 14 days. Trees were then treated with insecticides, imidacloprid, and carbaryl, to eliminate ACP, and the trees were kept free of ACP until the time of sampling. For branch analyses, all leaves for a given time point were detached via razor blades, flash-frozen, and held in liquid nitrogen during sampling. Petioles were removed and stored at −80 °C for *C*Las quantitation via qPCR. For metabolome analysis, ⅛ in (3.175 mm) diameter punches were dispensed into the wells of 96 well plates containing 500 µL 50% ethanol. Tissues were lysed by repeat freeze-thaw cycles between −80 °C and room temperature. The resulting extracts were filtered through 0.22 um filter plates via centrifugation before analysis via LC-MS. Leaf punches were collected in the same manner for the single-leaf assay. Collection of plant material comply with relevant institutional, national, and international guidelines and legislation

#### Field collected tissues

Stems, roots, and leaves from 50 trees (n=150) were collected from 5 different citrus Florida citrus groves in 2016 ^34^. In 2017, stems, roots, and leaves were collected from 80 trees located in 7 different orchards (n=240). Each tree was divided into 4 quadrants (North, South, East, and West), and stems with attached leaves were collected from each of the quadrants and pooled. Topsoil from two sides of the tree and approximately 1.5 feet away from the base of the trunk near the irrigation line was removed, and the feeder roots near the irrigation line were sampled, shaken to remove soil, and sealed in a plastic bag. Gloves were changed, and clippers and shovels were sterilized with 30% household bleach between each tree that was sampled. All samples were immediately placed on ice for transit to the laboratory, where they were placed at 4°C and processed within 24 hours.

### MS data acquisition

The tissue extracts were prepared in 100% ethanol, spiked with 1µM sulfadimethoxine internal standard, and analyzed with UltiMate 3000 UPLC system (Thermo Scientific) using a Kinetex^TM^ 1.7 µm C18 reversed-phase UHPLC column (50 X 2.1 mm) and Maxis Q-TOF mass spectrometer (Bruker Daltonics) equipped with ESI source. The column was equilibrated with 2% solvent B (98% acetonitrile, 0.1% formic acid in LC-MS grade water with solvent A as 0.1% formic acid in water), followed by a linear gradient from 2% B to 10% B in 0.2 min and then to 100% B at 12 min, held at 100% B for 2 min. Following each run, the column was equilibrated at 2% B for 1 min at a flow rate of 0.5 mL/min. MS spectra were acquired in positive ion mode in the range of 80-2000 m/z. A mixture of 10 µg/mL of each sulfamethazine, sulfamethizole, sulfachloropyridazine sulfadimethoxine, amitriptyline, and coumarin-314 was run at the beginning and the end of each batch (one 96-well plate). An external calibration with ESI-L Low Concentration Tuning Mix (Agilent Technologies) was performed prior to data collection, and internal calibrant Hexakis(1H,1H,3H-tertrafluoropropoxy)phosphazene was used throughout the runs. The capillary voltage of 4500 V, nebulizer gas pressure (nitrogen) of 1.4 bar, ion source temperature of 180 °C, and dry gas flow of 4 L/min, were used. For acquiring MS/MS fragmentation, the 7 most intense ions per MS^1^ were selected. A stepping function was used to fragment ions at 50%, 100%, 150%, and 200% of the CID, with a timing of 25% for each step.

Similarly, basic stepping of collision RF of 250 to 1500 Vpp with a timing of 25% for each step and transfer time stepping of 50, 75, 100, and 150 µs with a timing of 25% for each step was employed. MS/MS active exclusion parameter was set to 5 and released after 30 seconds. The mass of internal calibrant was excluded from the MS/MS list using a mass range of *m/z* 921.5–924.5. The data were deposited in the MassIVE online repository and are available under the IDs: MSV000082967 (2D leaf mapping); MSV000082962 (3D branch mapping); MSV000082963 and MSV000085416 (field study).

### MS data analysis

The collected HPLC-MS raw data files were first converted from Bruker’s *d* to mzXML format and then processed with the open-source MZmine2 software [https://www.ncbi.nlm.nih.gov/pubmed/20650010?dopt=Abstract]. crop filtering with a retention time (RT) range of 0 to 14 min chromatograms. Mass detection was performed with a signal threshold of 1E3 and a 0.04-s minimum peak width. The mass tolerance was set to 20 ppm, and the maximum allowed retention time deviation was set to 5 s. For chromatographic deconvolution, the local minimum search algorithm with a 30% chromatographic threshold, minimum RT range of .6 sec, minimum relative height of 1%, minimum absolute height of 5E2, the minimum ratio of peak top/edge 2, and peak duration range of 0.04 - min was used. After isotope peak removal, the peak lists of all samples were aligned within the corresponding retention time and mass tolerances. Gap filling was performed on the aligned peak list using the peak finder module with 1% intensity, 10-ppm *m/z* tolerance, and 0.05-min RT tolerance, respectively. After the creation and export of a feature matrix containing the feature retention times, exact mass, and peak areas of the corresponding extracted ion chromatograms, the sample metadata was added to the feature matrix metadata of the samples.

All of the peaks that were present in any of the blanks with a signal-to-noise ratio (S/N) below 3:1 were removed from the final feature table.

### Data pretreatment and statistical analysis

The data pretreatment and following statistical analysis were carried out with the MetaboAnalyst platform ^55^. The feature tables generated with MZmine were filtered to remove features with near-constant, very small values and values with low repeatability using the interquartile range (IQR) estimate. A detailed description of the methodology is given in ^56^. The samples were normalized using quantile normalization. The data were further scaled by mean centering and divided by standard deviation for each feature.

Principal component analysis (PCA) and partial least-squares discriminant analysis (PLS-DA) ^57^ were used to explore and visualize variance within data and differences among experimental categories. Random forests (RF) ^58^ supervised analysis was used to further verify the validity of determined discriminating features.

### Molecular networking

The molecular network was created using the online workflow at GNPS platform (gnps.ucsd.edu) ^17,18^. The data were clustered with MS-Cluster with a parent mass tolerance of 0.1 Da and an MS/MS fragment ion tolerance of 0.1 Da to create consensus spectra. The consensus spectra that contained less than 3 spectra were discarded. A network was then created where edges were filtered to have a cosine score above 0.65 and more than 4 matched peaks. The edges between two nodes were kept in the network if and only if each of the nodes appeared in each other’s respective top 10 most similar nodes. The spectra in the network were then searched against GNPS’s spectral libraries. All matches kept between network spectra and library spectra were required to have a score above 0.7 and at least 5 matched peaks. The molecular networks and the parameters used are available at the links below: 2D leaf mapping: https://gnps.ucsd.edu/ProteoSAFe/status.jsp?task=105598d2d782412ca0c988bfe933c032 3D branch mapping: https://gnps.ucsd.edu/ProteoSAFe/status.jsp?task=e2f1a1e367a7450fb940fe7cda04dc19 Field study: https://gnps.ucsd.edu/ProteoSAFe/status.jsp?task=2b0b1e554bc14d1fba321257b0ffc827 Field study, feature-based molecular network ^59^: https://gnps.ucsd.edu/ProteoSAFe/status.jsp?task=4ac8aff08af5433292143047c8cc5e90

### 2D/3D visualization

The procedure for the creation and visualization of 3D models is described in detail previously ^13^. Briefly, the 2D images and 3D model of the sampled plants were created and the coordinates for sampled spots were selected according to described protocol ^13^. The abundances of detected metabolites were normalized and autoscaled ^55^, and the coordinates for spots corresponding to each sample were inserted into the feature tables along with the spot size. The 2D or 3D models were drag-and-dropped into the ‘ili website in the browser (https://ili.embl.de/), followed by the feature table with coordinates. All figures were generated with the “Jet” color map. Spot size, opacity, and border opacity were adjusted for optimal visualization.

### Synthesis of feruloyl putrescine hydrochloride

As the feruloyl putrescine was not available commercially, we have synthesized this compound for further compound identification validation and disc assay testing. *N*-Boc putrescine (1 equiv, 4.25 mmol, 0.91 mL) and ferulic acid (1.06 equiv, 4.5 mmol, 880 mg) were combined in anhydrous CH_2_Cl_2_ (32.5 mL) with stirring and the solution was cooled to 0 °C. Then, a solution of N,N′-dicyclohexylcarbodiimide (DCC, 1.7 equiv, 7.2 mmol, 1.49 g) in anhydrous CH_2_Cl_2_ was added dropwise. The reaction mixture was allowed to warm to room temperature and stirred at for 2 days. Then, the mixture was filtered to remove precipitated dicyclohexylurea and the filtrate was concentrated *in vacuo*. The crude residue was purified over silica gel using 2-4% MeOH in CH_2_Cl_2_ to afford *N*-Boc feruloyl putrescine as a light yellow solid in 78% yield (1.29g). For Boc deprotection, trifluoroacetic acid (TFA, 15 mL) was added to a stirred solution of *N*-Boc feruloyl putrescine in CH_2_Cl_2_ (75 mL) under an inert atmosphere of Ar. The reaction was allowed to stir at room temperature for 40 min, then the solvent was removed *in vacuo.* The residue was dissolved in methanol (15 mL) with HCl (25 mL) and evaporated *in vacuo* to yield feruloyl putrescine hydrochloride as a light yellow solid (75% overall yield). ^1^H and ^13^C NMR data were consistent with those previously reported ^60^. HRMS (ESI) exact mass calculated for [M+H]^+^ (C_14_H_21_N_2_O_3_) is m/z 265.1547.

### Simulations

Simulations were performed using the *C*Las Ishi-1 M15 metabolic model^9^. Standard biomass constraints were maintained to predict the overall *C*Las growth rate. All model simulations were performed using the Gurobi Optimizer v.5.6.3 solver (Gurobi Optimization) in the COBRA toolbox ^61^ for MATLAB (MathWorks). We simulated the maximal growth rate of *C*Las using flux-balance analysis. The main metabolic compounds affecting *C*Las growth were identified using the metabolome data, that was anthranilate, ferulate, glutamate, N-acetyl-L-ornithine, ornithine, putrescine, tyrosine, and vanillin and sensitivity analysis, looking for metabolite-specific growth responses was performed by varying uptake rates among 1×10^-12^, 1×10^-10^, 1×10^-8^, 1×10^-6^, 1×10^-4^, 1×10^-2^, 1×10^-1^, 1×10^0^, 1×10^1^, 1×10^2^, 1×10^3^. Additionally, we performed a sensitivity analysis, which deployed a phenotypic phase plane that facilitates the observation of effects on the *C*Las growth by varying a particular constraint, in this case putrescine and ferulic acid. Predicted growth rates were compared with experimental results.

### Microbial culture assays

Citrus metabolites were assayed using a previously developed disc-diffusion assay for *L. crescens* ^27^. Briefly, *L. crescens* liquid cultures were incorporated to a soft agar (0.8%) overlay and applied to a 20 mL solid agar (1.5%). *L. crescens* strain BT-1] was maintained and grown exclusively on the previously described bBM7 + 1.0 methyl-β-cyclodextrin. To these overlaid plates were applied autoclave-sterilized 6 mm paper discs (Whatman, NJ, USA), previously loaded with 35 μL of a given metabolite solution and dried in a sterile biosafety cabinet. Once discs were applied, plates were sealed and stored upside down in a 28 ℃ incubator for 6 days to allow a clear zone of inhibition development for measurement. 35 mg/mL solutions of putrescine (Fisher Scientific, Waltham, MA, USA), and feruloylputrescine (synthesis described above) were prepared using sterile water.

### The efficacy of bioflavonoids and ferulic acid *in vitro C*Las-hairy root assay

The in vitro *C*Las-hairy roots assay was performed according to the previously described protocol ^28^. *C*Las-citrus hairy roots were generated using *C*Las-infected citrus plant tissues, and the diagnosis of *C*Las was confirmed by quantitative PCR (qPCR). *C*Las-hairy roots were surface sterilized, and ∼100 mg was transferred into multi-well plates containing Gamborg’s B-5 medium with 1% sucrose. Different concentrations of bioflavonoids (Horbaach, https://horbaach.com/products/citrus-bioflavonoids-complex-1500mg-300-vegetarian-caplets) and ferulic acid (Fisher Scientific, Catalog No. ICN10168505): 125, 250, 500, and 1000 ppm/mL, were added, vacuum infiltrated and incubated on a rotator shaker at 50 rpm in the dark at 25°C for 72 h. The experiments were carried out with six biological replicates, positive control (oxytetracycline hydrochloride), untreated *C*Las hairy roots, and an equal concentration of ethanol solvent used to dissolve the bioflavonoids and ferulic acid as negative controls. After the treatments, tissue samples were treated with PMAxx dye (propidium monoazide, Biotium, Fremont, CA) to inactivate dead *C*Las bacterial DNA. Further, total DNA was extracted, and viable bacterial titer was estimated by qPCR analysis using primers specific to the *C*Las gene encoding the Ribonucleotide reductase β-subunit (nrdB, RNR-F/RNR-R) ^62^ and the relative *C*Las titers were estimated and plotted relative to untreated using the 2−ΔΔCt method. After normalization of target Ct with an endogenous reference gene (Ct’) glyceraldehyde3-phosphate dehydrogenase 2 (GAPC2)^63^ to correct for DNA template concentration differences among the samples, it was plotted relative to untreated controls.

### Primers used in this study

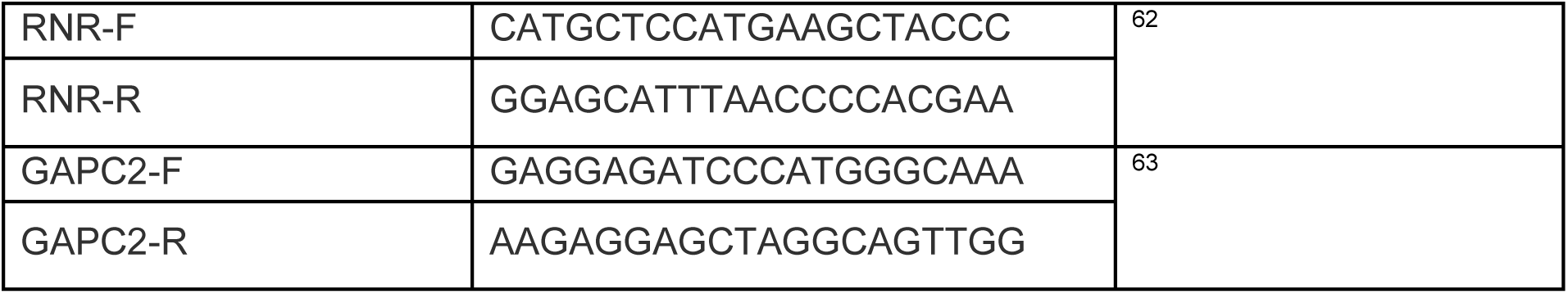

## Acknowledgements

This work was partly supported by USDA NIFA grant no. 2017-70016-26053; USDA NIFA grants 2021-70029-36056, and Texas A&M AgriLife Insect-vector disease seed grants to K.M.

## Author notes

Alexander A. Aksenov

Present address: Department of Chemistry, University of Connecticut, Storrs, CT, USA

Emily C. Gentry

Present address: **Department of Chemistry**, Virginia Tech, Blacksburg, VA

Cristal Zuniga

Present address: Department of Biology, San Diego State University, San Diego, CA, USA

Nichole Ginnan

Present address: One Health Microbiome Center, Huck Institutes of the Life Sciences, Pennsylvania State University, University Park, PA, USA

## Contributions

CR, PR, PCD created the idea for the work.

NG, AB, GM, PR collected plant tissues

NG, AB extracted plant tissues for acquisition of mass spectrometry data

AA acquired mass spectrometry data

AA, AM analyzed mass spectrometry data

GM conducted plants growth, infection by CLas, PCR diagnostics

AM created 3D models

AB, AM picked coordinates on plant models

AA performed statistical analysis

EG performed organic synthesis

CZ, KZ performed and analyzed model simulations AB performed disk assays

MR, KM conducted hairy roots assay AA wrote the manuscript

CR, AB, PR, NG, AL edited the manuscript

## Data availability statement

The data were deposited in the MassIVE online repository and are available below links:

1- https://massive.ucsd.edu/ProteoSAFe/dataset.jsp?task=bc1261c22e6c49d1b4f6490c7414845b

2- https://massive.ucsd.edu/ProteoSAFe/dataset.jsp?task=2008873282ac4beda2a008ada2917fa4

3- https://massive.ucsd.edu/ProteoSAFe/dataset.jsp?task=d4416036d13141b9b3008c7712395534

4- https://massive.ucsd.edu/ProteoSAFe/dataset.jsp?task=7db2bd0c5a0941f28286cabdc407a17f

5- https://massive.ucsd.edu/ProteoSAFe/dataset.jsp?task=25598a865760458fbc45a3f579c8479c

