## Supplementary information for "Spatial chemistry of citrus reveals molecules bactericidal to *Candidatus* Liberibacter asiaticus"

Supplemental

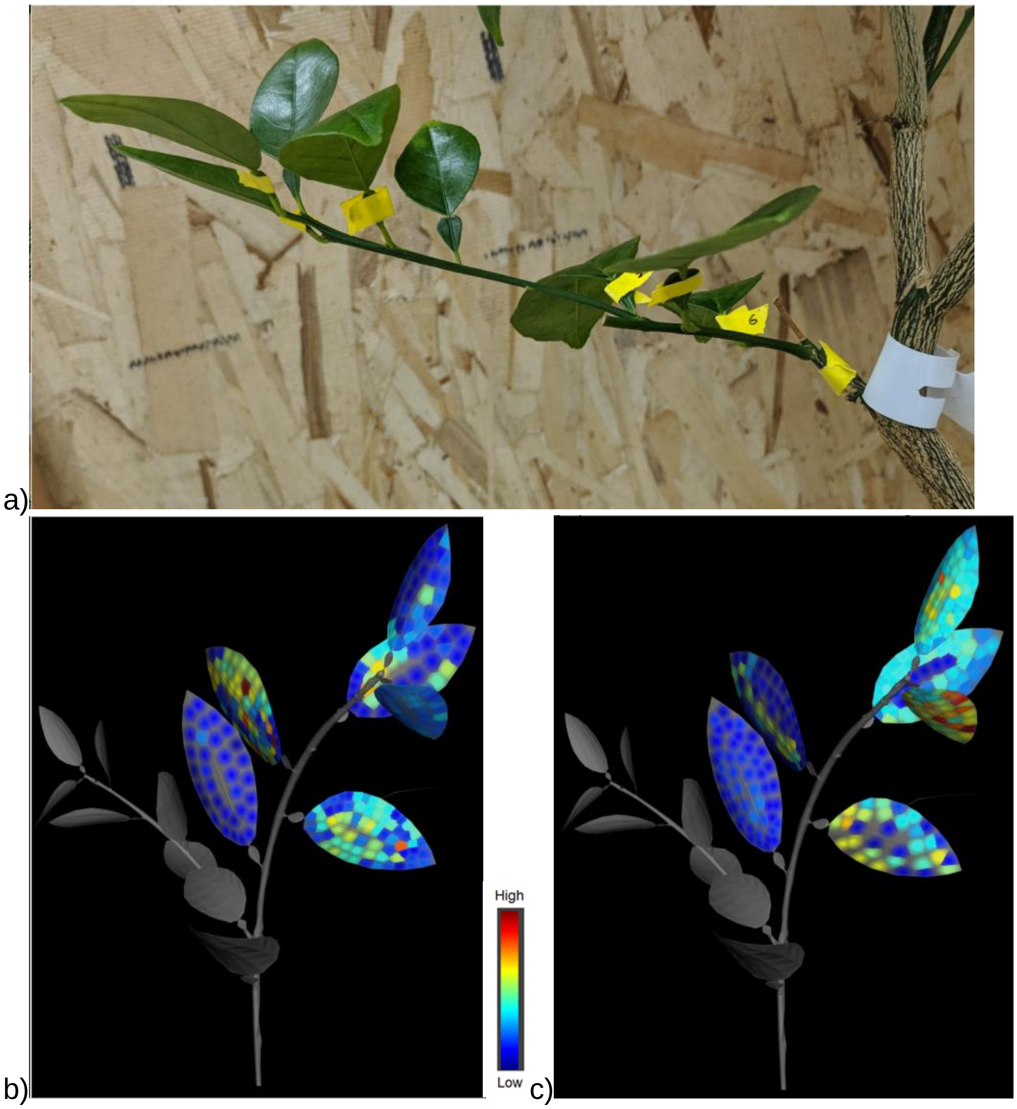


**Figure S1.** 3D branch mapping. a) The actual branch of a healthy sweet orange (Bridal) tree used for 3D mapping. Yellow tape is placed on the six leaves that were sampled for mapping. b) 3D map of rendered branch with abundances of 4',5,6,7-tetramethoxyflavone (scutellarein tetramethyl ether, m/z 343.1176 (rt 317.95)) mapped for the healthy and c) diseased trees' branches. The distribution of scutellarein tetramethyl ether was primarily affected by leaf age where it was detected at low abundance in older leaves and higher abundance in younger leaves at the apex of the branch.

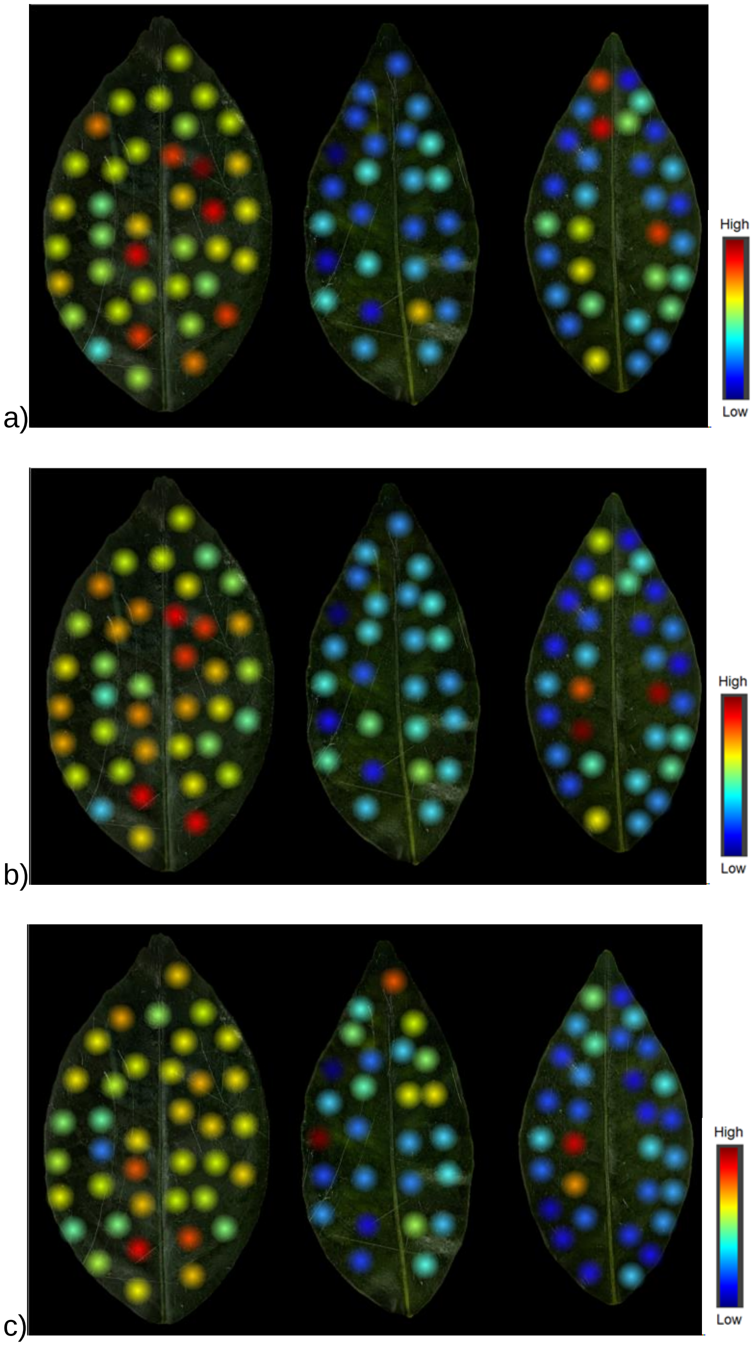


**Figure S2**. 2D map of: a) f) 4',5,6,7-Tetramethoxyflavone (m/z 343.1183, rt 317.59 sec) b) Tangeretin (m/z 373.1291, rt 341.57 sec) c) Neodiosmin (m/z 609.1824, rt 168.64 sec). 2D maps show from left to right: healthy, infected symptomatic, infected asymptomatic leaves.

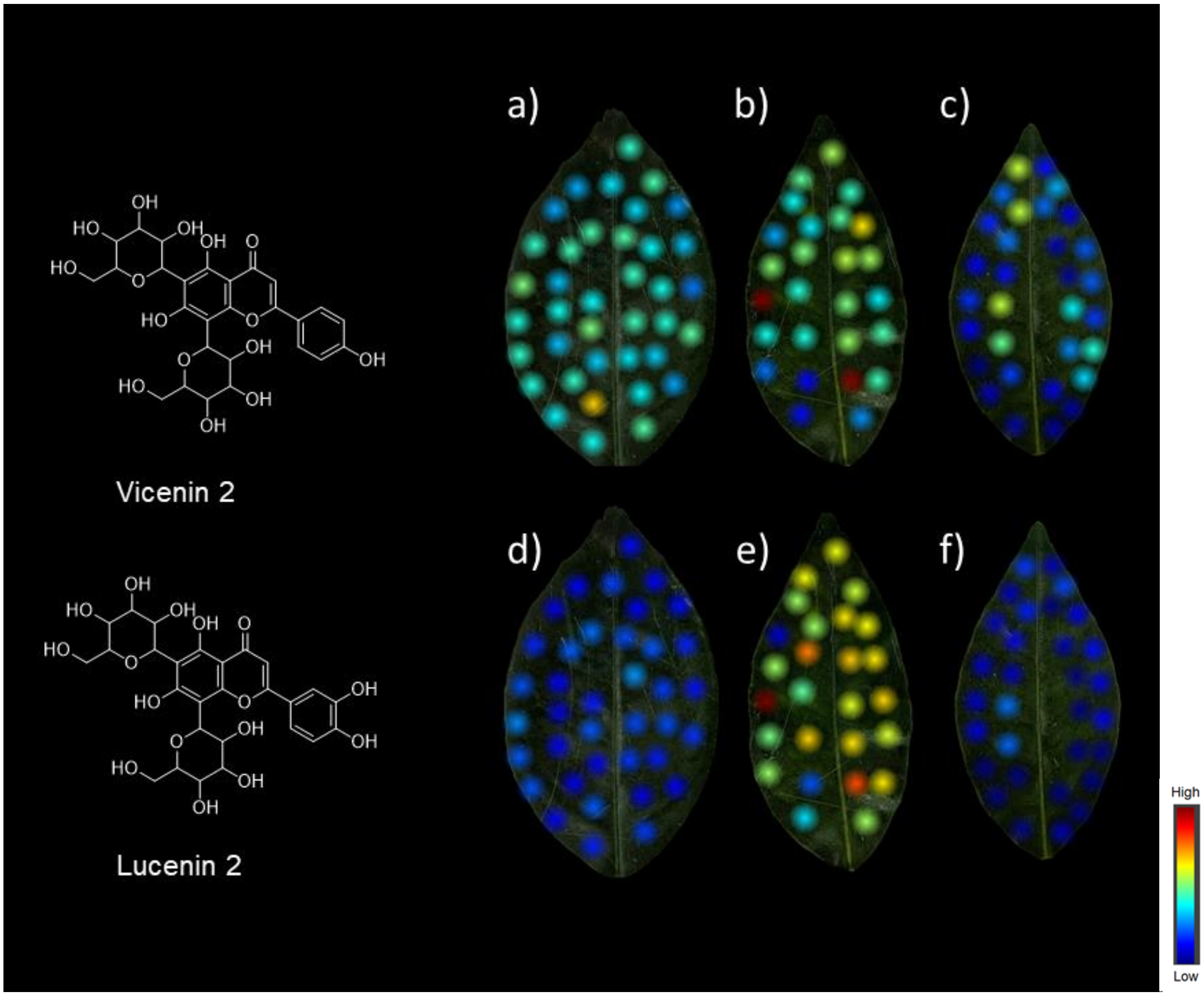


**Figure S3.** Vicenin 2 distribution in: a) Healthy b) Infected symptomatic c) Infected asymptomatic leaves; Lucenin 2 distribution in: d) Healthy e) Infected symptomatic f) Infected asymptomatic leaves.

a) b)

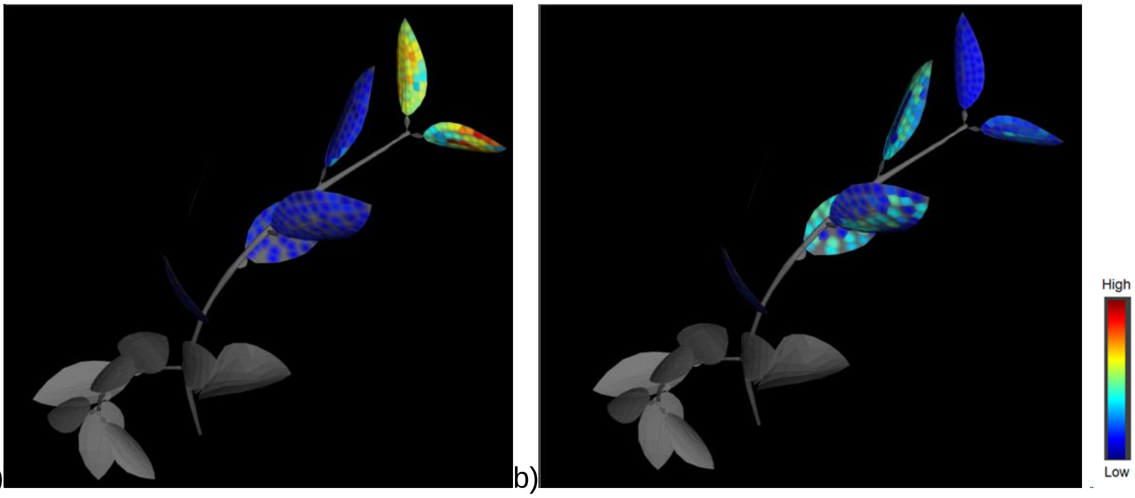


**Figure S4**. 3D map of: a) Feruloylputrescine, m/z 265.154 (rt 81.02 sec), infected plant b) Ferulic acid - H2O, m/z 177.0539 (rt 91.03 sec), infected plant

a) b)

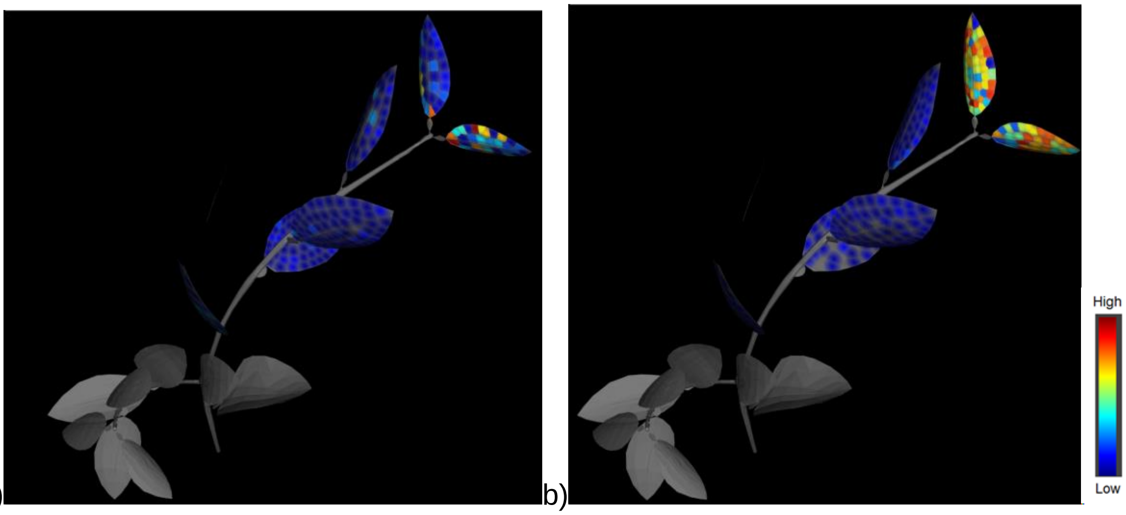


**Figure S5**. 3D map of: a) Hydroxycinnamic acid, m/z 165.0539 (rt 92.3 sec), healthy plant d) Hydroxycinnamic acid, m/z 165.0539 (rt 92.3 sec), infected plant
